# Efficient isolation of specific genomic regions retaining molecular interactions by the iChIP system using recombinant exogenous DNA-binding proteins

**DOI:** 10.1101/006080

**Authors:** Toshitsugu Fujita, Hodaka Fujii

## Abstract

**Background:** Comprehensive understanding of mechanisms of genome functions requires identification of molecules interacting with genomic regions of interest *in vivo*. We have developed the insertional chromatin immunoprecipitatin (iChIP) technology to isolate specific genomic regions retaining molecular interactions and identify their associated molecules. iChIP consists of locus-tagging and affinity purification. The recognition sequences of an exogenous DNA-binding protein such as LexA are inserted into a genomic region of interest in the cell to be analyzed. The exogenous DNA-binding protein fused with a tag(s) is expressed in the cell and the target genomic region is purified with antibody against the tag(s). In this study, we developed the iChIP system using recombinant DNA-binding proteins to make iChIP more straightforward.

**Results:** In this system, recombinant 3xFNLDD-D (r3xFNLDD-D) consisting of the 3xFLAG-tag, a nuclear localization signal, the DNA-binding domain plus the dimerization domain of the LexA protein, and the Dock-tag is used for isolation of specific genomic regions. 3xFNLDD-D was expressed using a silkworm-baculovirus expression system and purified by affinity purification. iChIP using r3xFNLDD-D could efficiently isolate the single-copy chicken *Pax5* (c*Pax5*) locus, in which LexA binding elements were inserted, with negligible contamination of other genomic regions. In addition, we could detect RNA associated with the c*Pax5* locus using this form of the iChIP system combined with RT-PCR.

**Conclusions:** The iChIP system using r3xFNLDD-D can isolate specific genomic regions retaining molecular interactions without expression of the exogenous DNA-binding protein in the cell to be analyzed. iChIP using r3xFNLDD-D would be more straightforward and useful for analysis of specific genomic regions to elucidate their functions.

## Background

Genome functions are mediated by various molecular complexes in the context of chromatin [1]. Comprehensive understanding of mechanisms of genome functions requires identification of molecules interacting with genomic regions of interest *in vivo*. To this end, we recently developed the locus-specific chromatin immunoprecipitation (ChIP) technologies consisting of insertional ChIP (iChIP) [2–5] and engineered DNA-binding molecule-mediated ChIP (enChIP) [6, 7] to isolate genomic regions of interest retaining molecular interactions. The functions of the genomic regions can be comprehensively understood by analysis of DNA, RNA, proteins, or other molecules interacting with the genomic regions.

In principle, iChIP is based on the locus-tagging strategy to isolate specific genomic regions using exogenous DNA-binding molecules. In contrast, enChIP is based on recognition of endogenous DNA sequences by engineered DNA-binding molecules such as TAL proteins and the CRISPR system. The scheme of iChIP is as follows: (i) The recognition sequences of an exogenous DNA-binding protein such as a bacterial protein, LexA, are inserted into the genomic region of interest in the cell to be analyzed. (ii) The DNA-binding domain (DB) of the exogenous DNA-binding protein is fused with a tag(s) and a nuclear localization signal(s) (NLS) and expressed in the cell to be analyzed. (iii) The resultant cell is stimulated and crosslinked with formaldehyde or other crosslinkers, if necessary. (iv) The cell is lysed, and the chromatin DNA is fragmented by sonication or enzymatic digestion. (v) The complexes including the exogenous DB are immunoprecipitated with antibody (Ab) against the tag(s). (vi) The isolated complexes which retain molecular interactions are reverse crosslinked, if necessary, and subsequent purification of DNA, RNA, proteins, or other molecules allows their identification and characterization. We successfully identified proteins and RNA components of an insulator, which functions as boundaries of chromatin domains [8], by using iChIP combined with mass spectrometry (iChIP-MS) or RT-PCR (iChIP-RT-PCR) [2]. Thus, iChIP is a useful technology for elucidation of molecular mechanisms of genome functions.

We recently developed 3xFNLDD, the second-generation tagged LexA DB consisting of 3xFLAG-tags, an NLS, and DB plus the dimerization domain of LexA, to utilize in iChIP [4]. 3xFNLDD is expressed in the cell to be analyzed for binding to the inserted LexA BE and subsequent purification of target genomic regions in the iChIP technology. If target genomic regions inserted with LexA BE can be pulled down using recombinant 3xFNLDD conjugated to Ab against the tags (Figure 1), expression of 3xFNLDD in the cell to be analyzed would not be necessary. In addition, it is not necessary to consider unexpected side effects of expression of 3xFNLDD on cell behavior, if any, making the procedure more straightforward.

**Figure 1.**
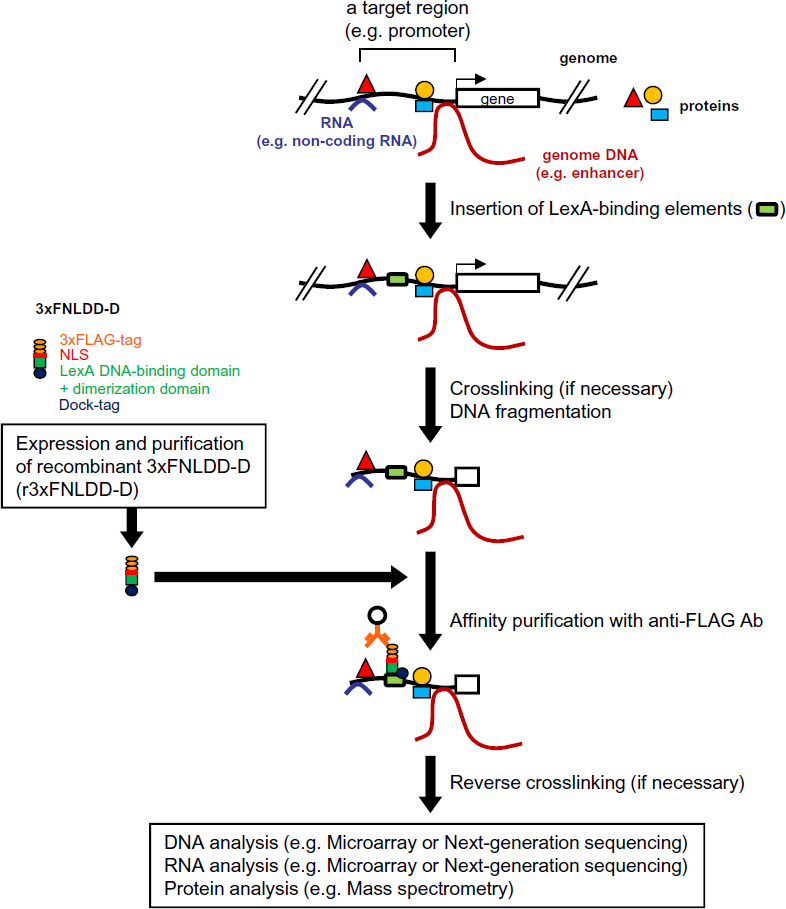
Scheme of iChIP using r3xFNLDD-D. 3xFNLDD-D consisting of 3xFLAG-tags, a nuclear localization signal (NLS), the DNA-binding domain (DB) plus the dimerization domain of the LexA protein, and Dock-tag, is expressed and purified. The recognition sequences of the LexA protein (LexA BE) are inserted into a genomic region of interest, usually by homologous recombination, in the cell to be analyzed. The resultant cell is stimulated and crosslinked with formaldehyde or other crosslinkers, if necessary. The cell is lysed, and the genomic DNA is fragmented. The target genomic region is affinity purified with r3xFNLDD-D conjugated with anti-FLAG antibody (Ab). After revers crosslinking, if necessary, purification of the chromatin components (DNA, RNA, proteins, other molecules) allows their identification and characterization.

In this study, we developed the iChIP system using the recombinant C-terminally Dock-tagged 3xFNLDD (3xFNLDD-D). Recombinant 3xFNLDD-D (r3xFNLDD-D) was expressed using a silkworm-baculovirus expression system and purified by affinity purification. iChIP using r3xFNLDD-D could effectively isolate the single-copy chicken *Pax5* (c*Pax5*) locus from a chicken B cell line, DT40. In addition, we could detect RNA associated with the c*Pax5* locus using this form of the iChIP system combined with RT-PCR. Thus, iChIP using r3xFNLDD-D would be more straightforward and useful to isolate specific genomic regions for their biochemical analysis.

## Results and Discussion

### Expression and purification of 3xFNLDD-D

For preparation of the purified r3xFNLDD-D, we utilized a silkworm-baculovirus expression system [9]. In this system, 3xFNLDD-D was expressed in a silkworm pupa by infection of baculoviruses expressing 3xFNLDD-D. The expressed protein was purified from the pupal homogenates using Dock Catch Resin, which specifically bound to Dock-tag in a calcium-dependent manner [9]. As shown in Figure 2A, SDS-PAGE followed by Coomassie Brilliant Blue (CBB) staining detected a single protein band at 35 kDa in the elution fraction. This protein was confirmed as 3xFNLDD-D by immunoblot analysis with anti-Dock Ab (Figure 2B). Thus, 3xFNLDD-D could be expressed in a silkworm pupa and purified without visible degradation.

**Figure 2.**
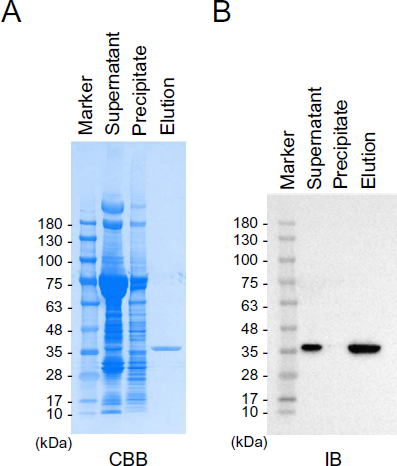
Expression and purification of r3xFNLDD-D. **(A)** Coomassie Brilliant Blue (CBB) staining. The purified proteins were subjected to SDS-PAGE and CBB staining. **(B)** Immunoblot analysis (IB). The purified proteins were subjected to SDS-PAGE and IB with anti-Dock Ab. Supernatant: the supernatant prepared from the silkworm pupal homogenates. Precipitant: the insoluble precipitate prepared from the silkworm pupal homogenates. Elution: the eluate after affinity purification with Dock Catch Resin.

### Efficient isolation of a target genomic region by iChIP using r3xFNLDD-D

Next, we examined whether the purified r3xFNLDD-D could be utilized for isolation of genomic regions of interest from vertebrate cells. To this end, we used the chicken DT40-devivative cell line, DT40#205-2, in which 8 x repeats of LexA BE were inserted 0.3 kbp upstream of the exon 1A of the single-copy endogenous c*Pax5* gene [10] (Figure 3A). The crosslinked chromatin prepared from the cell line was subjected to iChIP using r3xFNLDD-D as shown in Figure 1. After purification of the immunoprecipitated genomic DNA, the yield of the c*Pax5* 1A promoter region was evaluated by detection of the LexA BE site (LexA BE) and the region 0.2 kbp upstream of LexA BE (i.e., 0.7 kbp upstream of the transcription start site (TSS) of c*Pax5* exon 1A) (-0.7k) by real-time PCR (Figure 3A). As shown in Figure 3B, the yield of LexA BE and -0.7k were more than 20% and 5% of input, respectively, when 10 µg of each r3xFNLDD-D and anti-FLAG Ab were used. In contrast, the yield of the genomic region 10 kbp upstream of the TSS of the exon 1A (-10k) was less than 0.01%. These results suggested that r3xFNLDD-D can bind to LexA BE even in the crosslinked chromatin and iChIP using r3xFNLDD-D is able to specifically purify target genomic regions. The specific isolation of the c*Pax5* 1A promoter region was completely blocked when we inhibited binding of r3xFNLDD-D to anti-FLAG Ab with excessive amounts of 3xFLAG peptide (Figure 3C). The c*Pax5* 1A promoter region was not isolated when parental DT40 was used instead of DT40#205-2 (Figure 3C). These results clearly demonstrated that isolation of target genomic regions is mediated by binding of LexA BE with r3xFNLDD-D captured by anti-FLAG Ab.

**Figure 3.**
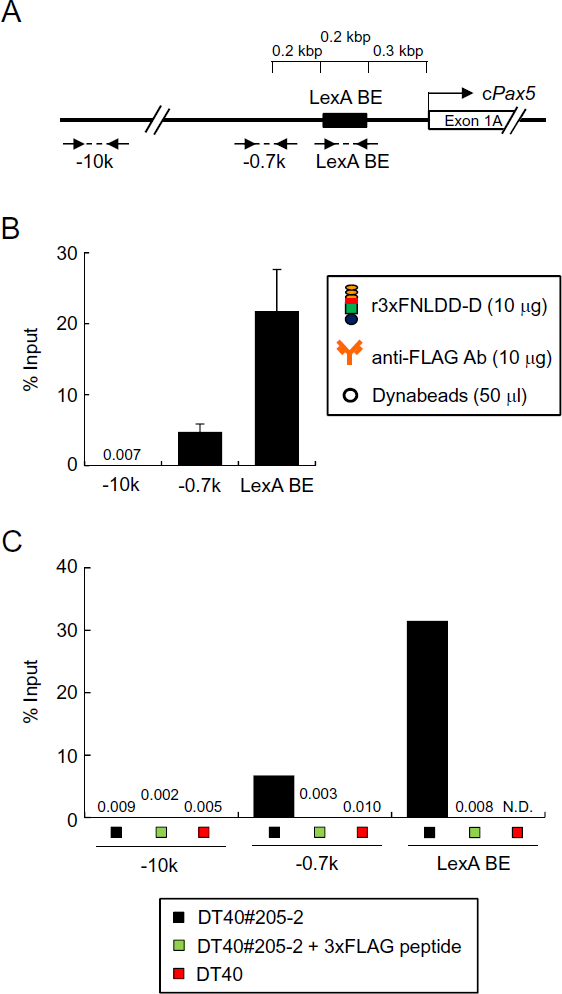
Isolation of the c*Pax5* 1A promoter region by iChIP using r3xFNLDD-D. **(A)** Scheme of the LexA BE-inserted c*Pax5* 1A promoter region with primer positions. The positions of PCR primers with distances from the transcription start site (TSS) are indicated. **(B)** The results of iChIP using 10 μg of r3xFNLDD-D. % of input is shown (mean +/-SEM, n = 3). **(C)** Specific isolation of the target genomic region by iChIP using r3xFNLDD-D. N.D.: not detected.

### Optimization of iChIP using r3xFNLDD-D

Next, we titrated amounts of r3xFNLDD-D and anti-FLAG Ab to optimize the system (Figure 4A). The yield of the LexA BE site in the c*Pax5* 1A promoter was comparable when 0.5 - 10 μg of each r3xFNLDD-D and anti-FLAG Ab was used with chromatin prepared from 1 × 10^7^ DT40#205-2 cells. In contrast, use of 0.01 - 0.1 μg of each protein showed lower yield, suggesting that 0.5 μg of each r3xFNLDD-D and anti-FLAG Ab are sufficient for 1 × 10^7^ cells. The yield of iChIP using 0.5 μg of r3xFNLDD-D was 20% of input for LexA BE and less than 0.01% for -10k, which is comparable with that using 10 μg of r3xFNLDD-D (Figures 3B and 4B). We also examined whether it would be possible to purify the c*Pax5* 1A promoter region using r3xFNLDD-D with Dock Catch Resin, which bound to the C-terminal Dock-tag of r3xFNLDD-D. As shown in Supplemental Figure 1, 2% of input of the LexA BE site could be isolated with negligible contamination of -10k, indicating that the C-terminal Dock-tag of r3xFNLDD-D can also be utilized for iChIP, although the yield was much lower than that using anti-FLAG Ab.

**Figure 4.**
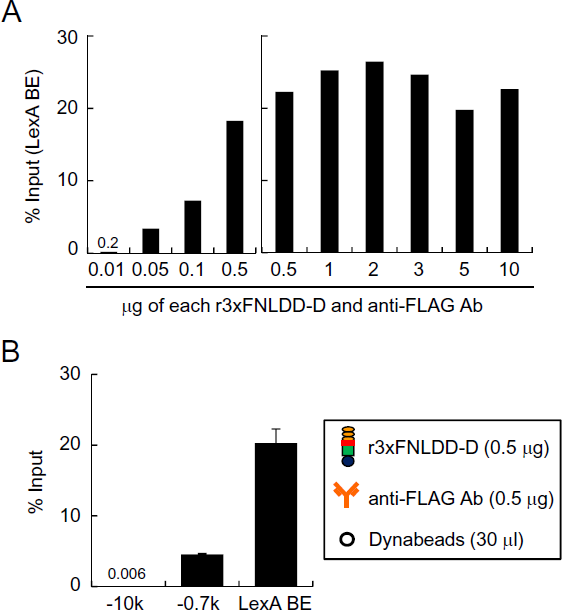
Optimization of iChIP using r3xFNLDD-D for isolation of the c*Pax5* 1A promoter region. **(A)** Titration of r3xFNLDD-D and anti-FLAG Ab. **(B)** The results of iChIP using 0.5 μg of each r3xFNLDD-D and anti-FLAG Ab. % of input is shown. The error bar represents the range of duplicate experiments.

### Isolation of RNA associated with the c*Pax5* locus by iChIP using r3xFNLDD-D

Next, we examined whether iChIP using r3xFNLDD-D could be utilized to isolate and identify molecules interacting with genomic regions of interest in cells. To this end, we attempted to isolate the c*Pax5* locus including the exon 1A region by iChIP using r3xFNLDD-D and detect the nascent RNA transcribed from the TSS of the c*Pax5* exon 1A by RT-PCR (Figure 5A). The c*Pax5* exon 1A was transcribed even in the presence of LexA BE inserted in the 1A promoter region (Supplemental Figure 2). As shown in Figure 5B, iChIP using r3xFNLDD-D isolated the c*Pax5* exon 1A region but not the exon 3 region of the irrelevant c*AID* gene, which encodes an enzyme essential for B cell-specific immunoglobulin somatic hypermutation and class switch recombination [11]. After purification of the associated RNA, RT-PCR analysis detected RNA transcribed corresponding to the exon 1A of the c*Pax5* gene but not corresponding to the c*AID* gene in the iChIP sample (Figure 5C) (the full-length images with size markers are shown in Supplemental Figure S3). These results suggested that iChIP using r3xFNLDD-D is able to isolate specific genomic regions retaining molecules interacting with the genomic regions.

**Figure 5.**
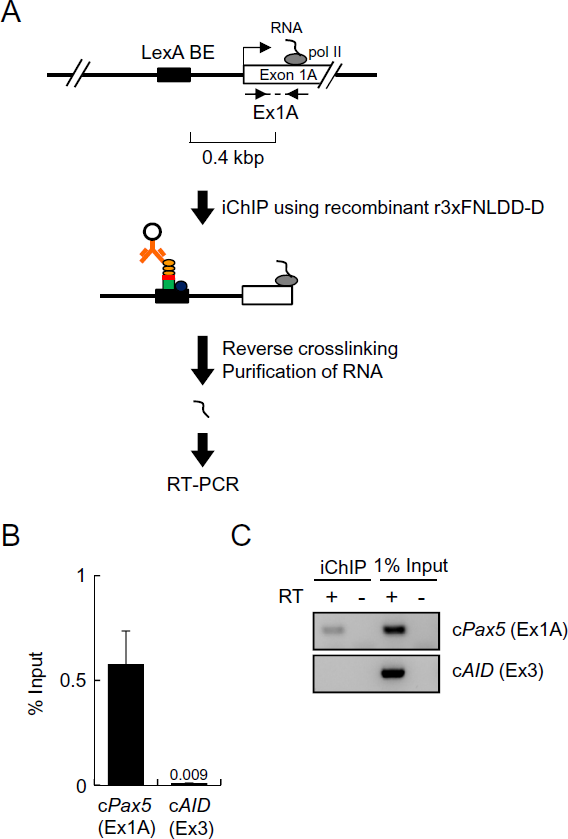
Detection of RNA associated with the c*Pax5* locus. **(A)** Scheme of iChIP. After isolation of the c*Pax5* locus by iChIP using r3xFNLDD-D, the nascent RNA transcribed on the exon 1A of the c*Pax5* gene was detected by RT-PCR. **(B)** The results of iChIP using 0.5 μg of each r3xFNLDD-D and anti-FLAG Ab. % of input is shown. The error bar represents the range of duplicate experiments. **(C)** Detection of RNA corresponding to the exon 1A of the c*Pax5* gene but not that corresponding to the exon 3 of the c*AID* gene by RT-PCR.

In this study, we applied RT-PCR to detection of RNA interacting with a genomic region of interest in cells. Next-generation sequencing or microarray analysis can be combined with iChIP using r3xFNLDD-D for non-biased identification of interacting RNA as well as DNA. Moreover, mass spectrometry can be combined for non-biased identification of interacting proteins.

Because iChIP using r3xFNLDD-D does not require expression of 3xFNLDD in cells, it is of great use in the iChIP analysis of primary cells isolated from organisms, especially higher eukaryotes such as mice. In the case of application of the standard iChIP technology to mice, it is time-consuming to establish mouse lines expressing 3xFNLDD in the cells to be analyzed as well as those possessing LexA BE in specific genomic regions. In this regard, iChIP using r3xFNLDD-D is able to skip the mating steps between mice expressing 3xFNLDD and those possessing LexA BE and the subsequent backcrossing steps, substantially accelerating iChIP analysis using organisms.

### Conclusions

In this study, we established the iChIP system using r3xFNLDD-D to make the iChIP technology much more straightforward. Using this system, we were able to isolate target genomic regions; % of input reached more than 20% for the c*Pax5* 1A promoter region. In addition, we could detect RNA associated with the c*Pax5* locus, suggesting that iChIP using r3xFNLDD-D can isolate target genomic regions retaining molecular interactions. Thus, iChIP using r3xFNLDD-D would be more straightforward and useful for analysis of specific genomic regions to elucidate their functions.

## Materials and Methods

### Expression and purification of r3xFNLDD-D

Expression of 3xFNLDD-D was performed using the silkworm-baculovirus expression system (ProCube) (Sysmex Corporation, http://procube.sysmex.co.jp/eng/) as described previously [9]. Briefly, the coding sequence of 3xFNLDD [4] was inserted into the transfer vector pM31a (Sysmex Corporation) and co-transfected with linearized genomic DNA of ABv baculovirus (*Bombyx mori* nucleopolyhedrovirus; CPd strain, Sysmex Corporation) into the *B. mori*-delived cell line, BmN, to generate the recombinant baculovirus. The generated baculovirus was infected into a silkworm pupa to express 3xFNLDD-D. The expressed 3xFNLDD-D was purified with Dock Catch Resin (Sysmex Corporation) as described previously [9]. The immunoblot analysis was performed with anti-Dock Ab (Sysmex Corporation).

### Cell lines

The chicken B cell line DT40 was provided by the RIKEN BioResource Center through the National Bio-Resource Project of the Ministry of Education, Science, Sports and Culture of Japan. DT40 and DT40#205-2, in which LexA BE was inserted in the 1A promoter region of the c*Pax5* gene (Fujita and Fujii, manuscript submitted), were maintained in RPMI-1640 (Sigma) with 4 mM glutamine, 10% (v/v) fetal bovine serum, 1% chicken serum, and 50 μM 2-mercaptoethanol at 39.5°C.

### Chromatin preparation

Cells (2 × 10^7^) were fixed with 1% formaldehyde at 37°C for 5 min. The chromatin fraction was extracted and fragmented (2 kbp-long on average) by sonication as described previously [12] except for using 800 µl of *in vitro* Modified Lysis Buffer 3 (10 mM Tris pH 8.0, 150 mM NaCl, 1 mM EDTA, 0.5 mM EGTA) and Ultrasonic disruptor UD-201 (TOMY SEIKO). After sonication, TritonX-100 was added to final concentration at 0.1%.

### iChIP using r3xFNLDD-D

The sonicated chromatin (400 µl) was pre-cleared with 0.01 - 10 μg of normal mouse IgG (Santa Cruz Biotechnology) conjugated to 30 - 50 μl of Dynabeads-Protein G (Invitrogen) and subsequently incubated with 0.01 - 10 μg of r3xFNLDD-D and anti-FLAG M2 Ab (Sigma) conjugated to 30 - 50 μl of Dynabeads-Protein G at 4°C for 20 h. 100 μg of 3xFLAG peptide was added to inhibit binding of r3xFNLDD-D to anti-FLAG Ab. The Dynabeads were washed four times with 1 ml of *in vitro* Wash Buffer (20 mM Tris pH 8.0, 150 mM NaCl, 2 mM EDTA, 0.1% TritonX-100) and once with 1 ml of TBS-IGEPAL-CA630 (50 mM Tris pH 7.5, 150 mM NaCl, 0.1% IGEPAL-CA630). The isolated chromatin complexes were eluted with 120 µl of Elution Buffer (500 µg/ml 3xFLAG peptide (Sigma-Aldrich), 50 mM Tris pH 7.5, 150 mM NaCl, 0.1% IGEPAL-CA630) at 37°C for 30 min. After reverse crosslinking at 65°C, DNA was purified with ChIP DNA Clean & Concentrator (Zymo Research) and used as template for real-time PCR with SYBR Select PCR system (Applied Biosystems) using the Applied Biosystems 7900HT Fast Real-Time PCR System. PCR cycles were as follows: heating at 50°C for 2 min followed by 95°C for 10 min; 40 cycles of 95°C for 15 sec and 60°C for 1 min. The primers used in this experiment are shown in Table 1.

**Table 1.**
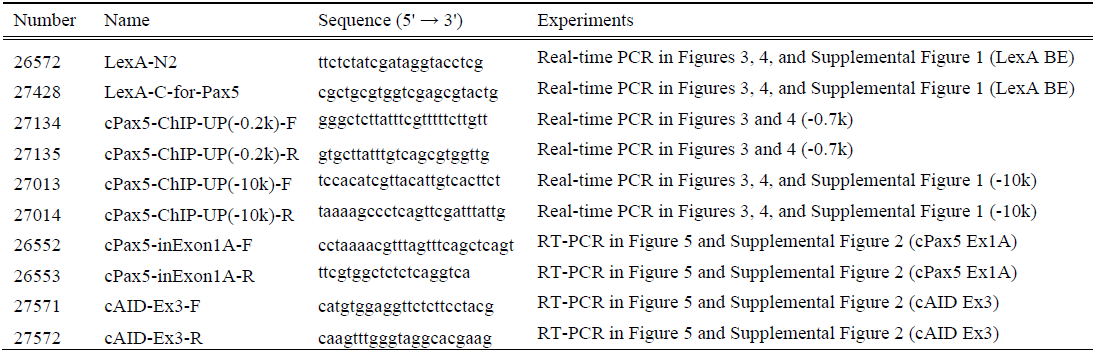
Primers used in this study.

### Isolation of interacting RNA and RT-PCR

Chromatin preparation and iChIP using recombinant r3xFNLDD-D were performed as described above except for addition of RNasin Plus RNase Inhibitor (Promega) in all buffers. After reverse crosslinking at 65°C, RNA was isolated with Isogen II (Nippon gene) combined with Direct-zol RNA Mini Prep (Zymo Research). The purified RNA was used as template for reverse transcription with ReverTra Ace qPCR RT Master Mix with gDNA Remover (Toyobo). The cDNA was used as template for PCR with AmpliTaq Gold 360 Master Mix (Applied Biosystems). PCR cycles were as follows: heating at 95°C for 10 min; 40 cycles of 95°C for 30 sec, 60°C for 30 sec, 72°C for 1 min; and the final extending 72°C for 2 min. The primers used in this experiment are shown in Table 1.

## Competing interests

Patents on iChIP are already registered (‘‘A method to isolate specific genomic regions’’, August 20, 2010, PCT/JP2010/064052; April 9, 2013, US patent : 8415098; November 22, 2013, Japanese patent : 5413924). This does not alter the authors’ adherence to all the BMC Molecular Biology policies on sharing data and materials.

## Authors’ contributions

T.F. and H.F. conceived this form of the iChIP technology, designed and performed experiments, and wrote the manuscript. H.F. directed and supervised the study.

## Acknowledgements

We thank F. Kitaura for technical assistance. This work was supported by Takeda Science Foundation (T.F.), the Uehara Memorial Foundation (H.F.), the Kurata Memorial Hitachi Science and Technology Foundation (T.F. and H.F.), Adaptable & Seamless Technology Transfer Program through Target-driven R&D (A-STEP) by the Japan Science and Technology Agency (JST) (#AS251Z01861Q) (H.F.), Grant-in-Aid for Young Scientists (B) (#25830131) (T.F.), “Transcription Cycle” (#25118512) (H.F.) from the Ministry of Education, Culture, Sports, Science and Technology of Japan.

